# Open Iris - An Open Source Framework for Video-Based Eye-Tracking Research and Development

**DOI:** 10.1101/2024.02.27.582401

**Authors:** Roksana Sadeghi, Ryan Ressmeyer, Jacob Yates, Jorge Otero-Millan

## Abstract

Eye-tracking is an essential tool in many fields, yet existing solutions are often limited for customized applications due to cost or lack of flexibility. We present OpenIris, an adaptable and user-friendly open-source framework for video-based eye-tracking. OpenIris is developed in C# with modular design that allows further extension and customization through plugins for different hardware systems, tracking, and calibration pipelines. It can be remotely controlled via a network interface from other devices or programs. Eye movements can be recorded online from camera stream or offline post-processing recorded videos. Example plugins have been developed to track eye motion in 3-D, including torsion. Currently implemented binocular pupil tracking pipelines can achieve frame rates of more than 500Hz. With the OpenIris framework, we aim to fill a gap in the research tools available for high-precision and high-speed eye-tracking, especially in environments that require custom solutions that are not currently well-served by commercial eye-trackers.

**CCS CONCEPTS:**

- **Applied computing** → Life and medical sciences.

## 1 INTRODUCTION

Eye tracking technology is used in an ever-growing list of fields. In basic science research, the study of eye movements contributes to a comprehensive understanding of various facets of human brain function, such as vision [Gibaldi and Banks, 2019], vestibular processing [Cullen, 2023], [Otero-Millan and Kheradmand, 2016], attention [Kelley et al., 2008], motor learning [Sedaghat-Nejad and Shadmehr, 2021], and emotions [Zhang et al., 2022]. In clinical settings, eye tracking systems play a crucial role in medical diagnosis across various disciplines including neurology [Leigh and Zee, 2015], ophthalmology [Grillini et al., 2020], otology [Korda et al., 2021], psychology [Walker-Smith et al., 2013], and psychiatry [Takahashi et al., 2021]. The field of human-computer interfaces extensively relies on eye trackers to enhance communication between humans and computers/robots [Adhanom et al., 2023], [Duvinage et al., 2011]. Additionally, marketing research harnesses the power of eye tracking systems to objectively evaluate the efficacy of different advertisements or designs to capture and retain attention. [Casado-Aranda et al., 2023]

There is a wide variety of eye tracking technologies developed over the last decades. Each with unique features and compromises tailored to specific applications. Some are small and can be head mounted [Huang et al., 2024], others are large taking up almost entire room [Moon et al., 2024], some are expensive and high frequency, while others are affordable and low frequency. Eye tracking systems differ significantly in the range of eye movement amplitudes they can optimally measure with some of them only being able to measure large eye movements, others only small movements, and very few systems performing well over small and large movements [Stevenson et al., 2010]. Recently, video-based eye tracker systems, in particular, have become the most common choice in head mounted or desk mounted configurations because they offer valuable features at a potentially lower cost than other types of eye trackers. A high-quality video-based eye tracker can deliver adequate precision, high-speed real-time tracking, and a wide detection range for eye movements from less than 0.5 degrees to more than 20 degrees [Stevenson et al., 2010]. However, options for commercially available video-based eye trackers are limited, expensive and are often closed systems and may lack flexibility and customizability for specialized applications (for example EyeLink ®, Tobii, and Eyegaze Inc.).

As an alternative to commercial eye trackers, many open-source eye tracking solutions have surfaced addressing either the customization or cost-effectiveness needed in specialized research applications. These solutions are software based but often include recommendations for commercial hardware components so an eye tracker can be built and used with the software. The hardware components necessary for video-based eye tracking are typically: a video camera with a lens, an infrared (IR) illuminator, an IR filter, and a computer to acquire and process the video. Table 1 compares the features of different open-source software solutions available at the moment for video-based eye trackers. These solutions vary greatly in the set of features they provide. Only some of can collect data at a high frequency (500Hz or more). Many are implemented for a particular hardware configuration and will not be compatible with other hardware. Some only process the camera images online and do not allow for the recording of videos and most do not allow post-processing videos. These two features are especially useful in the development of new tracking algorithms and pipelines. Finally, not all of them include a graphical user interface. As shown in Table 1, a comprehensive platform offering high-performance, customizable features for different applications, along with a user-friendly graphical interface is still missing.

**Table 1:** Comparing Available open-source remote eye tracking systems.

| Name | Year | Frame Rate (Hz) | Hardware Compatibility | Post-Process | Binocular | Graphical User-Interface | Software Language |
| --- | --- | --- | --- | --- | --- | --- | --- |
| ITU Gaze Tracker [ITUGazeTracker, 1999] | 2010 | 30 | Yes | No | No | Yes | C# |
| GazeParser [Sogo, 2013] | 2013 | 500 | Yes | NA | Yes | No | Python |
| Pupil [Kassner et al., 2014] | 2014 | 22 | No | No | Yes | NA | Python, C |
| Oculomatic [Zimmermann et al., 2016] | 2016 | 600 | Yes | No | No | Yes | C++ |
| EyeRecToo [Santini et al., 2017] | 2017 | 125 | Yes | No | Yes | Yes | C++ |
| openEyeTrack [Casas and Chandrasekaran, 2019] | 2019 | 715 | No | NA | NA | NA | C++ |
| DeepVOG [Yiu et al., 2019] | 2019 | 130 | No | No | No | No | Python |
| RemoteEye [Hosp et al., 2020] | 2020 | 500 | No | Yes | Yes | Yes | C/C++ |
| EyeLoop [Arvin et al., 2021] | 2021 | 1000 | Yes | No | No | Yes | Python |
| Using DeepLabCut [Zdarsky et al., 2021] | 2021 | 30 | No | No | Yes | NA | Python |
| A low-cost, high-performance video-based binocular eye tracker [Ivanchenko et al., 2021] | 2021 | 395 | No | No | Yes | NA | C++ |
| OpenIris | 2024 | 500 | Yes | Yes | Yes | Yes | C# |

Here we introduce an open-source, extendable, high-performance, and user-friendly platform for video-based eye tracker systems that meets the reliability and speed requirements for research. Our framework, OpenIris, implements many features that are commonly required for eye-tracking systems, including multi-threading, data buffering, video and data recording, logging, graphical interfacing, and network communication. What separates OpenIris from other solutions is its modular design. By leveraging a plugin-based architecture, OpenIris enables developers to extend it to work with any hardware system and develop and deploy custom hardware configurations, tracking pipelines, and calibration routines. Here, we describe the structure and features of OpenIris and explain the options for developing new customizable plugins. At the end, we report examples of the performance of OpenIris with sample plugins for pupil and torsional tracking.

## 2 OPENIRIS FEATURES

OpenIris is written in the C# programming language, and relies on OpenCV [OpenCV, 2024] and its wrapper for C# emguC to achieve real-time binocular performance exceeding 500 frames per second. C# provides a good balance between ease of development and performance. OpenIris currently targets .Net Framework 4.8 and is developed as a Windows application, but future versions will be upgraded to target newer versions of .Net to become cross-platform and run under Windows, Linux and macOS. OpenIris is available at: https://github.com/ocular-motor-lab/OpenIris

### 2.1 Core Features

OpenIris already implements a set of core features common to most eye-tracking systems. This architecture allows plugin developers to focus on the image processing techniques for tracking or the hardware integration without the need to implement the often more tedious requirements involving user interface, real-time multithreading, data storage, error handling, and configuration.

#### 2.1.1 User Interface

The graphical interface of OpenIris, shown in Figure 1, is designed for user-friendly navigation and control. It features a setup tab that displays eye images in real-time, in addition to adjustable settings defined by the tracking pipeline plugin being used. The viewer tab presents eye data traces alongside corresponding videos, offering a comprehensive view of the tracking output. The calibration tab facilitates straightforward monitoring and adjustment of calibration process, with an interface defined by the calibration plugin in use. A log tab is also available to display errors or messages. Finally, a configuration window comprises three columns that allow users to set parameters for OpenIris in general or for each of the three main plugins.

**Figure 1.**
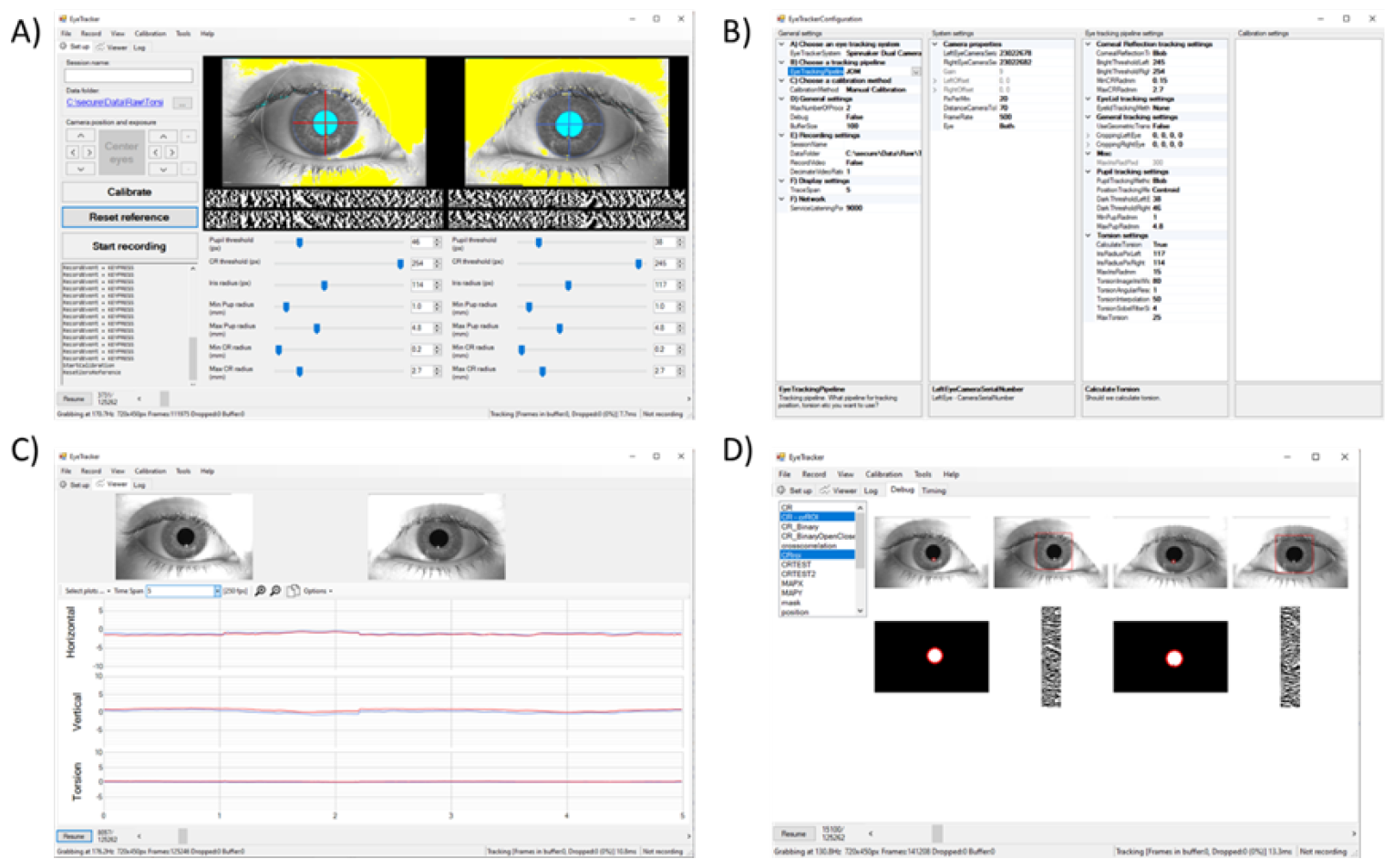
Example print-screen of the OpenIris a) main window setting tab with adjustable parameters, b) configuration window to select the plugins, c) view tab that shows online traces, and d) debug tab that can be used for debugging pipeline performance.

For advanced users and developers, a debug mode can be enabled which offers an additional tab with a detailed view of tracked objects at different processing stages. This feature aids in debugging the tracking pipeline as well as providing insights into the system’s performance. The plugin developer does not need to implement the user interface elements, they just need to indicate that a particular image during any intermediate processing step should be added to the debug tab.

#### 2.1.2 Remote Control

In addition to its graphical user interface, OpenIris offers alternative methods of interaction. OpenIris can be utilized purely as a library, providing an API to incorporate it in any other software. Alternatively, OpenIris features network interfaces to remotely communicate with other pieces of software via UDP, TCP, or HTTP protocols. This enables synchronization with other programs written in different languages such as Python or Matlab running on different devices. This feature allows remote control of the eye-tracking system, providing flexibility for experiment designs, such as running experiments in complete darkness or implementing gaze contingent displays.

#### 2.1.3 Offline / Batch Analysis

OpenIris offers offline and batch analysis features, allowing users to reprocess recorded videos with different parameters to refine eye traces. To reprocess multiple videos, the user can adjust the parameters for each video and process all videos in a batch.

#### 2.1.4 Recording

Data recording includes two primary outputs: a text file containing the results of frame-by-frame tracking, and, optionally, the raw videos obtained from the cameras. The videos are recorded in raw format to allow for their post processing without any loss, which is fundamental in the development and optimization of new tracking methods. This option requires storage solutions capable of the required transfer rates. OpenIris will inform the user if not all the frames are being recorded but it will continue to track.

#### 2.1.5 Buffering and Multithreading

OpenIris natively implements an image buffer which connects the video acquisition and image processing plugins. Due to the variation in processing time caused by non-constant-time algorithms or CPU usage by other processes, some frames may take substantially longer to process than the sampling rate of the eye cameras. OpenIris can often eliminate frame drops – which are caused by these spikes in processing time – by storing up to a set number of unprocessed frames in a first-in-first-out buffer. By reducing the size of this image buffer, it is possible to prioritize real-time latency at the risk of dropping frames.

Additionally, OpenIris runs plugin-defined image processing pipelines in contained threads, which allows for otherwise single-threaded algorithms to leverage multiple processing cores without special development. The number of threads can be easily adjusted by the user using the configuration window.

#### 2.1.5 Configuration

OpenIris stores all configurations, including system and plugin settings, as a serializable class. This architecture allows new plugins to define settings simply by defining their type and default value as properties in a class that inherits from a base settings class. OpenIris then automatically loads and interprets these settings at run time using reflection, provides a user interface to edit their values, and saves the settings in a text file. All of these features are handled by core structure, eliminating complexity during plugin development.

### 2.2 Plugins

The modular structure of OpenIris provides the flexibility of building three plugins (Figure 2a) that can be loaded into OpenIris as independent libraries without the need to rebuild the entire platform, and it can be interchanged at run time. First, it allows for plugins for different video camera hardware and configurations, accepting adjustable camera features such as frame rate, frame size, orientation, etc., depending on the features available in the user’s camera settings. Second, plugins to add or modify different tracking methods for measuring pupil movements, iris rotations, reflection movements, or even head movements. A third type of plugin is used for calibration methods.

**Figure 2.**
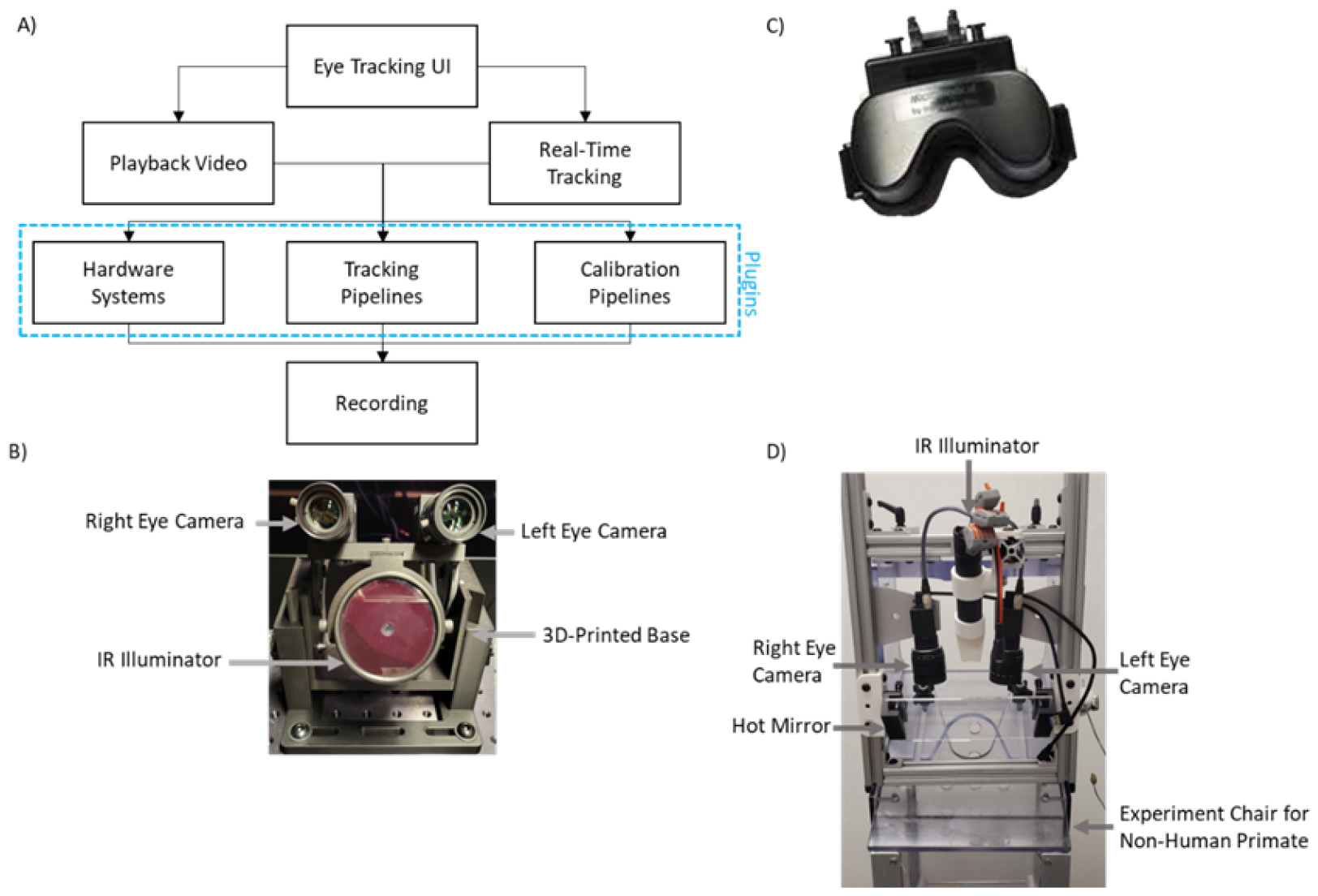
a) Simplified structure of OpenIris program, b) Example video-based eye tracking setup that used in OpenIris desktop system with two cameras and one IR illuminator c) Micromedical head-mounted eye tracking system by Intracoustics [Intracoustics, 2000] and d) setup used for non-human primate studies.

#### 2.2.1 Camera hardware systems

The main function of OpenIris’ hardware system plugins is to configure and initialize user-selected camera(s) to capture raw images. Because different cameras have unique settings associated with their characteristics, developers have the flexibility to initialize, start, and capture images from their chosen camera(s) by providing specific code snippets. In addition, the design of the OpenIris user interface enables users to create options for camera(s) settings to be dynamically changed during online tracking.

As an example of a hardware system for a binocular eye tracker, we have developed a plugin to work with two Blackfly S USB3 04S2M Flir cameras (with a resolution of 720 × 540 and a frequency of up to 500 frames per second) [Blackfly-S-USB3, 2024], each paired with a 50mm focal length lens and an IR filter. A 3D-printed base holds the cameras and an infrared illuminator. (Figure 2b)

Within the OpenIris systems plugin, we take advantage of the camera API to synchronize the capture of the two cameras via hardware connection. In this case, the left-eye camera is assigned as Primary, which sends a signal to the right eye camera to indicate when it should open the shutter and capture a new image. This ensures perfect temporal alignment of the two cameras. Additionally, within the plugin, the user interface was modified to add an option for setting the tracking frequency as a one-time adjustable parameter at the onset of the tracking process before camera initialization. Moreover, the gain of the cameras and region of interest is adjustable, allowing real-time modifications during tracking.

Figure 2b, c, and d display some of the different configurations that OpenIris has been adapted for. These include desktop systems with one or two cameras, head mounted goggles, and animal recording rigs.

#### 2.2.2 Tracking pipelines

The main function of tracking pipeline plugins is to process the images and extract from them relevant features about eye movements. Each camera frame enters the image processing pipeline, which may employ the image processing libraries, such as “OpenCV” [OpenCV, 2024] and its wrapper for C# emguC, for locating and extracting objects of interest. Similar to hardware systems settings, the OpenIris allows developers to add features to the user interface for adjusting image processing settings in real-time during tracking such as brightness thresholds, or minimum and maximum sizes for objects being tracked. (Table 2)

**Table 2:**
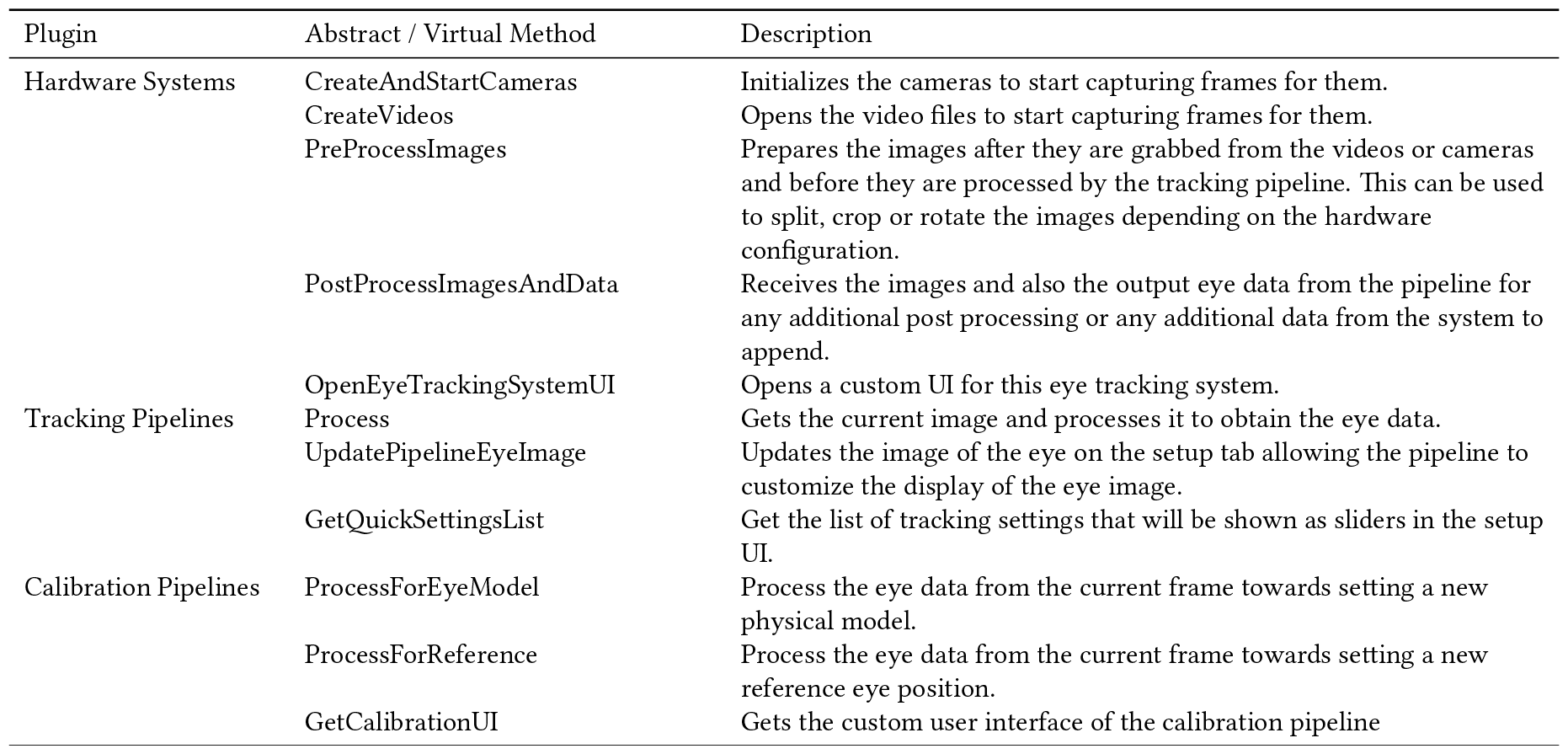
List of methods to be implemented by the plugins.

For instance, a tracking pipeline is already implemented within OpenIris to locate the pupil in an infrared camera frame using different methods including centroid, convex hull or ellipse fitting. In addition to the pupil tracking, the pipeline includes detecting corneal reflections (also known as glints) for pupil – CR method, and tracks the iris for torsional eye movements (Figure 3b,c). [Otero-Millan et al., 2015]

**Figure 3.**
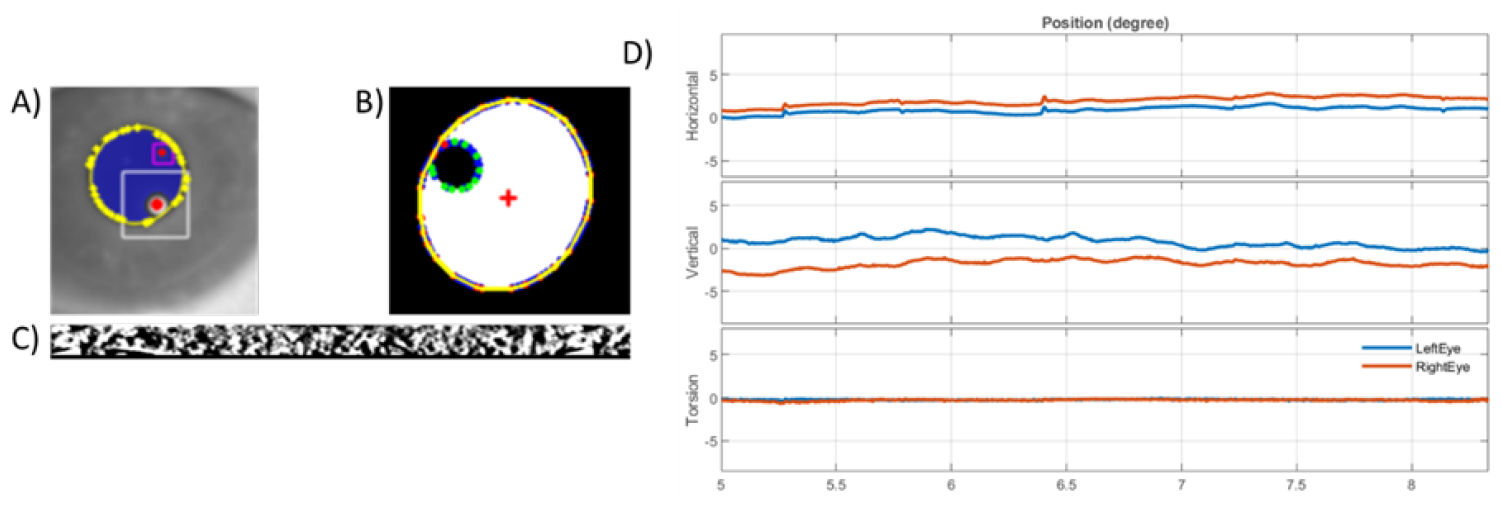
Example of detecting a) p1 and p4 shown in red circles for digital DPI, b) corneal reflection in green and pupil in yellow contour for pupil – CR method, and c) detecting iris pattern for tracking torsional eye movement. d) Example eye traces in degree for left eye in blue and right eye in red color.

An example of externally developed plugins is provided by the independent contributor (second author), who has developed a dual-Purkinje eye tracking pipeline for OpenIris that detects reflections of the illuminator from the cornea surface and the inner surface of the lens (Figure 3a). [Wu et al., 2023]

#### 2.2.3 Calibration pipelines

The main function of calibration pipeline plugins is to establish a mapping between the features extracted by the tracking pipeline and the actual motion of the eye. This mapping can be achieved either through image and geometric analysis or by analyzing behavior in interaction with a display. Initially, raw eye-tracking data is represented in units of pixel on the image. A calibration method translates these pixel values into degrees, corresponding to visual angles.

The calibration pipelines in OpenIris facilitate this conversion process by allowing users to define their own calibration methods. (Table 2) These methods can be based on established eye models or implemented through regression models that correlate known locations with corresponding eye positions. This modular approach empowers users to customize the calibration process to their specific experimental needs, ensuring accurate and meaningful conversion of pixel data to visual angles. The calibration pipeline implemented for behavioral calibration also includes network communication between a computer presenting the targets and a computer executing OpenIris to record the eye positions.

## 3 EXAMPLE USAGE

The goal of this article is to present the framework and its modular structure and not providing an extensive performance analysis because it ultimately depends on the specific plugins being used. As a reference we provide some example performance measurement for processing time and data quality, from tests conducted on a simple configuration for binocular pupil tracking utilizing two cameras and one illuminator. A participant looked at the central dot for a total of 6 minutes (with 20s intervals and 5s rests). OpenIris recorded the data at 500 Hz and was controlled within a Matlab R2021b program on a PC with processor of i9-11900 at 2.50 GHz, and installed RAM of 64.0 GB.

The study was approved by the University of California, Berkeley Institutions IRB and followed the tenets of the Declaration of Helsinki. The subject signed an informed consent form to participate after being informed of the goals and methods of the study.

### 3.1 Process Time and Data Quality

For images of size 720 × 540 pixels, using a maximum of five processing threads the baseline average time from a frame arriving at the computer to the completion of data processing, without any image processing, was 0.14 ± 0.18 ms (mean ± standard deviation). Introducing a simple pupil tracking, centroid method, the processing time was 0.79 ± 0.70 ms.

Using more advanced tracking methods for pupil detection, such as Ellipse fitting, the processing time increased to 2.50 ± 1.80 ms, tracking pupil and corneal reflection increased it to 3.70 ± 3.20 ms and tracking pupil and torsional eye movement to 9.70 ± 6.70 ms. The recordings included zero percent dropped frame in all conditions.

The saved video from one session was post-processed at 500 Hz with pupil detection (using ellipse fitting method) and torsion detection. Figure 3d shows an example horizontal, vertical and torsional eye positions. The root mean square eye velocity in horizontal (h) and vertical (v) directions with pupil were: h 18.55 deg/s, v 40.98 deg/s, and the root mean square eye velocity for torsion was: 19.99 deg/s.

## 4 SUMMARY

In conclusion, OpenIris is a user-friendly and adaptable tool for eye-tracking research. Its modular design allows easy customization, and key features like the remote control and batch analysis enhance its usability. Looking forward, expanding the tracking algorithms available and fostering collaborative development could further enhance its utility. OpenIris stands as a valuable and versatile resource for eye-tracking studies, balancing simplicity with the flexibility needed for various research applications.

## ACKNOWLEDGMENTS

This research was supported by the National Eye Institute NEI R00EY027846 and National Science Foundation NSF 2107049.

